# Region-specific *Per1* induction dissociates circadian and mood responses to light

**DOI:** 10.64898/2026.05.21.726853

**Authors:** Dan-Adrian Epuran, Jürgen A. Ripperger, Urs Albrecht

## Abstract

Light is a major environmental cue that synchronizes circadian rhythms and modulates affective behaviors. In mammals, photic information reaches multiple brain regions, including the suprachiasmatic nuclei (SCN), the master circadian pacemaker, and the lateral habenula (LHb), a structure implicated in mood-related behaviors such as despair. Both regions harbor the light-inducible clock gene *Per1*, which contributes to circadian phase shifting and behavioral regulation. Although light can simultaneously shift circadian phase and improve mood-related behaviors in mice and humans, it remains unclear whether these processes share a common mechanism. To address this question, we selectively deleted *Per1* in SCN or LHb neurons of *Per1*-floxed mice using region-specific viral Cre recombinase delivery. Loss of *Per1* in the SCN reduced light-induced phase advances at circadian time (CT) 22, whereas deletion in the LHb had no effect on phase shifting. In contrast, light-induced improvement of despair-like behavior was abolished by *Per1* deletion in the LHb but preserved in SCN-targeted knockouts. Furthermore, according to RT-PCR, *Per1*-induction relied on different promoter usage. These findings demonstrate that the mood-related effects of light are independent from its circadian phase-shifting properties, revealing distinct *Per1*-dependent mechanisms through which light regulates behavior. This dissociation suggests that light-based therapies could be optimized by targeting mood-regulating pathways without necessarily altering circadian activity cycles.

**Author summary:** The present study identifies *Per1* as a molecular node through which light exerts two distinct therapeutic actions: circadian phase resetting via the SCN and mood improvement via the LHb. This dissociation has major translational implications. It suggests that the antidepressant effects of light therapy—widely used in seasonal affective disorder (SAD) and increasingly in major depressive disorder (MDD)—do not require circadian phase shifting and instead rely on a *Per1*-dependent LHb pathway. Understanding this separation opens the door to more targeted, faster, and potentially more effective light-based interventions.

## Introduction

Most organisms on Earth are governed by a circadian clock system that aligns their physiology and behavior with the environmental light–dark cycle. In mammals, light serves as a potent synchronizing cue, transmitted by specialized retinal photoreceptors [1, 2] to multiple brain regions [3]. These include the suprachiasmatic nuclei (SCN), the master pacemaker of the circadian system, and the lateral habenula (LHb), a region implicated in the regulation of despair and despair-like behavior in both humans and mice. Light input entrains the molecular clocks within these neuronal populations, thereby coordinating central and peripheral oscillators with the prevailing photoperiod [4].

At the molecular level, circadian rhythmicity is generated by a transcription–translation feedback loop driven by the heterodimeric complex BMAL1 (Basic Helix-Loop-Helix ARNT-Like 1) and CLOCK (Circadian Locomotor Output Cycles Kaput). This complex activates transcription of its own negative regulators, the *Period* (*Per1/Per2*) and *Cryptochrome* (*Cry1/Cry2*) genes. The resulting feedback loop produces ∼24-hour oscillations in gene expression, influencing up to 20% of cellular transcripts in a tissue-specific manner (clock-controlled genes, CCGs) [5]. Disruption of core clock components, such as *Per1*, therefore leads to measurable alterations in circadian-regulated behaviors.

Light modulates circadian behavior by inducing phase shifts [6]. Nocturnal light exposure directly activates *Per* gene expression, thereby altering clock phase [7–10]. The direction of the shift depends on circadian timing: light at circadian time (CT) 14, corresponding to early active phase in mice, induces phase delays, whereas light at CT22, late in the active phase, produces phase advances [11].

Several brain regions, including the SCN and LHb, are directly or indirectly influenced by light [12–14]. Light signals processed in the SCN regulate the timing of locomotor activity onset [15], whereas light input to the LHb modulates despair-like behavior, as assessed by the forced-swim test (FST) [16]. In both cases, induction of *Per1* appears critical: loss of *Per1* reduces light-induced phase advances [10, 17], while loss of *Per1* in the LHb increases immobility in the FST following a brief light pulse [16]. Whether these two light-dependent processes are mechanistically linked or operate independently remains unclear.

This question has translational relevance. In humans, light facilitates adaptation to jet lag and exerts therapeutic effects in mood disorders such as seasonal affective disorder [18]. Moreover, the degree of circadian misalignment in major depressive disorder (MDD) correlates with symptom severity [19, 20]. Because behavioral phase shifts and mood improvements often occur concurrently following light treatment, it has been hypothesized that mood enhancement depends on light-induced circadian phase shifting [21].

To determine whether light-induced phase shifting and light-induced improvement of despair-like behavior are mechanistically linked or separable, we generated mice lacking *Per1* specifically in SCN or LHb neurons using viral delivery of a Cre-recombinase expression vector in *Per1*-floxed mice [16]. Mice lacking *Per1* in the SCN (SCN*^Per1^*^nKO^) exhibited reduced phase advances in response to light at CT22, whereas *Per1* deletion in the LHb (LHb*^Per1^*^nKO^) had no effect on phase shifting. Conversely, light-induced improvement in despair-like behavior at CT22 was preserved in SCN*^Per1^*^nKO^ animals but abolished in LHb*^Per1^*^nKO^ mice.

These findings indicate that the beneficial effects of light on despair-like behavior are independent of its phase-shifting effects on the circadian clock.

## Results

### Deletion of *Per1* in SCN neurons but not in the LHb shortens period

Previous studies have shown that a brief polychromatic light pulse (white light) delivered two hours before light onset induces *Per1* gene expression in both the SCN [9] and the LHb [16]. Whereas *Per1* induction in the SCN has been linked to phase advances in locomotor activity [10], *Per1* induction in the LHb has been associated with reduced despair-like behavior [16]. These findings raise the possibility that light-induced mood improvement may depend on circadian phase shifting, as suggested in humans [21].

To test this hypothesis, we generated mice lacking neuronal *Per1* either in the SCN (SCN*^Per1^*^nKO^) or in the LHb (LHb*^Per1^*^nKO^)(Fig 1A). *Per1* floxed (*Per1^fl/fl^*) mice received injections of adeno-associated virus (AAV) serotype 6 (targetting LHb) or 9 (targetting SCN) carrying either a control construct expressing EGFP under the synapsin promoter or the same construct additionally expressing iCre [16] (Fig 1B). These animals were subsequently assessed for circadian phase-shifting responses using wheel-ruinning activity and for despair-like behavior using the tail suspension test (TST). Daily wheel-running activity was comparable across all groups (Fig 1C). However, mice with *Per1* deletion specifically in SCN neurons (SCN*^Per1^*^nKO^ mice) exhibited a significalty shorter free-running period compared to both control and LHb*^Per1^*^nKO^ mice (Fig 1D), consistent with previous findings that loss of *Per1* in brain neurons shortens circadian period [22]. Thus, *Per1* deletion in SCN neurons – but not in LHb neurons – selectively shortens circadian period.

**Fig 1.**
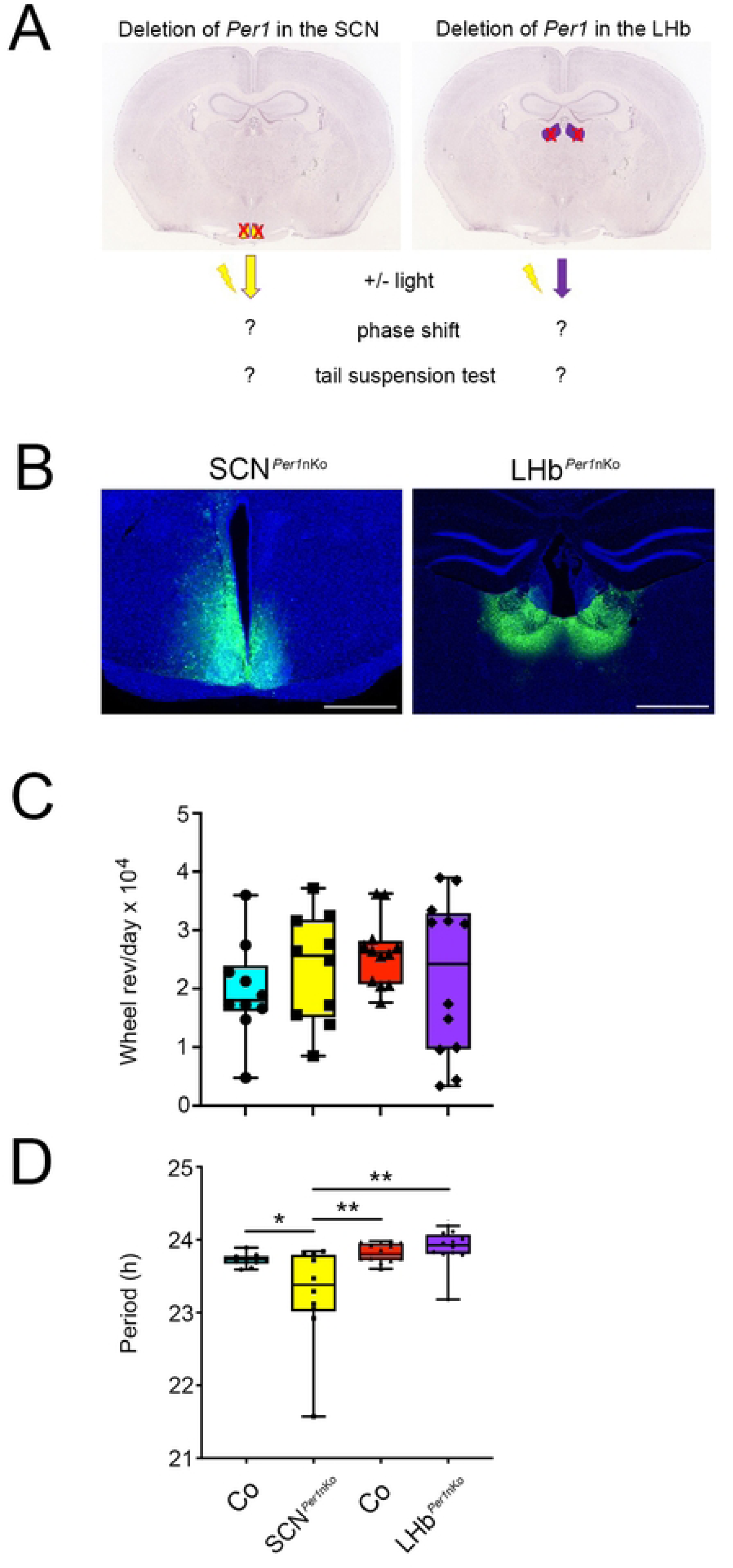
*Per1* deletion in neurons of the SCN but not LHb shortens period. (A) Scheme illustrating the experimental set up. Deletion of *Per1* in the SCN (left panel, yellow) and in the LHb (right panel, purple) is illustrated. Below the panels, the behavioral tests before and after a light pulse are indicated. (B) Photomicrographs of SCN and LHb after injection of virus; blue color represents nuclei (DAPI staining) and green shows injected virus expressing green fluorescent protein (GFP). Scale bar: 500 µm. (C) Total wheel-running activity of control mice and animals with a local deletion of *Per1* in neurons (*Per1*nKo) of the SCN (green and yellow bars) or LHb (red and purple bars). No significant differences were observed. Data are represented as mean ± SEM, n = 10-12, one-way ANOVA with Tukey’s multiple comparison test, p > 0.05. (D) Period in SCN*^Per1^*^nKo^ (yellow bar), LHb*^Per1^*^nKo^ (purple bar) and their corresponding control animals (Co, green and red bars) is shown. Period of SCN*^Per1^*^nKo^ mice is significantly shorter compared to the other experimental groups. Data are represented as mean ± SEM, n = 10-12, one-way ANOVA with Tukey’s multiple comparison test, *p < 0.05, **p < 0.01.

### Light-induced phase shifts require *Per1* in SCN neurons but not in LHb neurons

To determine how *Per1* deletion affects light-induced phase shifting, we used an Aschoff Type I protocol under constant darkness (DD) conditions [11]. The experimental design is shown in Figure 2A. Mice were maintained in DD with access to running wheels, and locomotor activity was recorded and displayed as double-plotted actograms (Fig 2B). After ∼10 days in DD, animals received a 15-minute light pulse (LP) at circadian time (CT) 10. Two weeks later they received an LP at CT14. After another two weeks a final LP at CT22 was applied (Fig. 2B, yellow stars). The selected time points correspond to regions of the mouse phase response curve (PRC): CT10 (no phase shift), CT14 (phase delay), and CT22 (phase advance) [11]. Following each LP, mice remained in DD, and activity onsets were marked before (blue lines) and after (red lines) the LP, and the displacement between these lines was used to quantify phase shifts. Quantitative analysis confirmed that LPs at CT10 produced no significant phase shifts in any group, consistent with the PRC (Fig 2C, left panel). LPs at CT14 induced phase delays in all groups, but delays were significantly larger in SCN*^Per1^*^nKO^ mice compared with all other genotypes (Fig 2C, middle panel). LPs at CT22 elicited phase advances in all groups except SCN*^Per1^*^nKO^ mice, which showed significantly reduced advances relative to controls (Co) and LHb*^Per1^*^nKO^ animals (Fig 2C, right panel).

**Fig 2.**
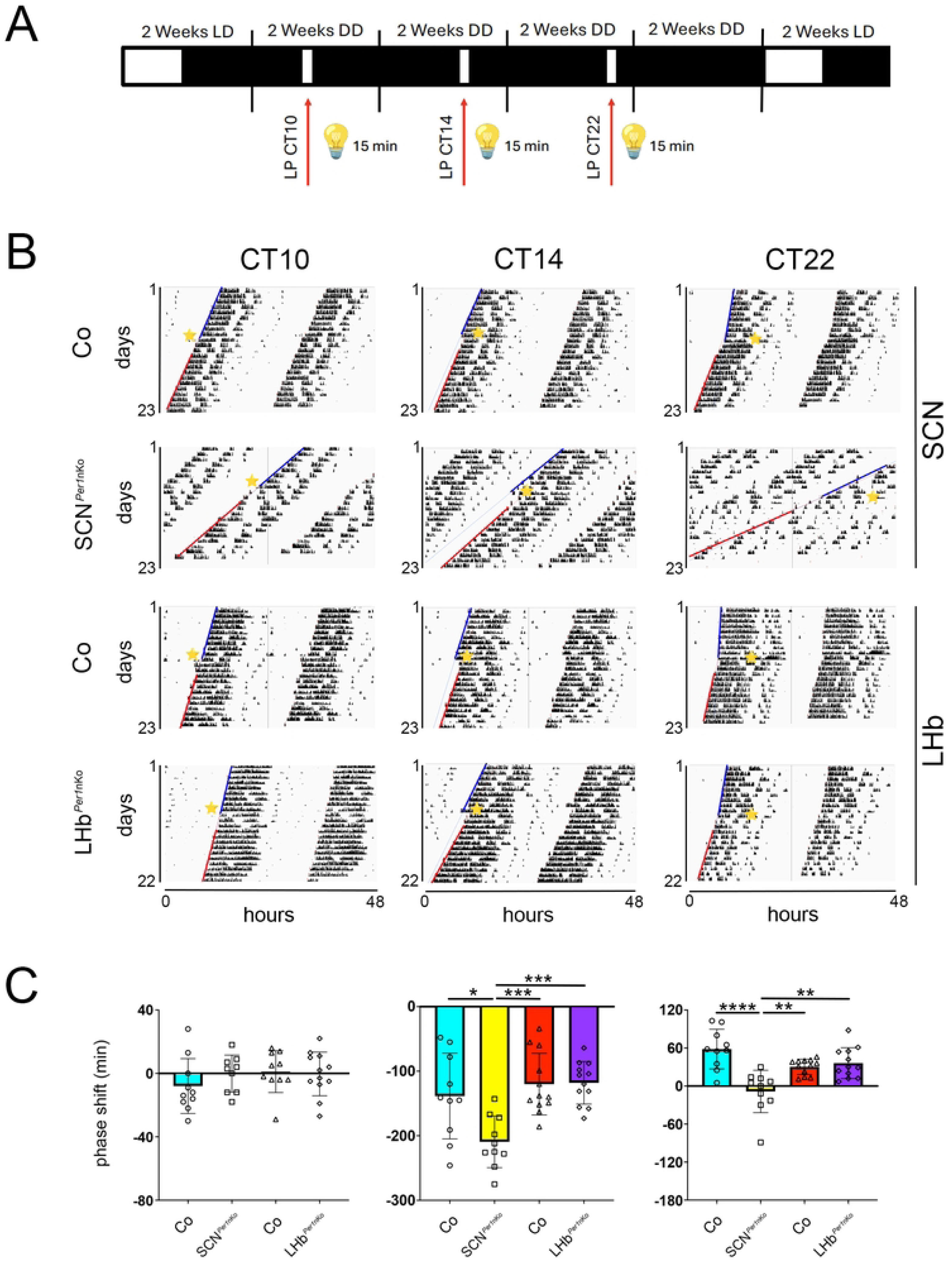
*Per1* deletion in neurons of the SCN but not the LHb affects phase shifts of the circadian clock. (**A**) Experimental design for application of 15-minute light pulses (LPs, 1000 LUX, red arrows) using the Aschoff type I protocol. Mice were kept in a light / dark cycle (LD) for 2 weeks and then they were released into constant darkness (DD) before receiving a LP at circadian time (CT) 10. Subsequently, the animals were kept for 2 weeks in DD before they received a LP at CT14 (early subjective night). The mice were again kept for 2 weeks in DD and then they received a LP at CT22 (late subjective night). Subsequently, the animals were kept again in DD for 2 weeks. During the whole experiment locomotor activity was recorded. White and black bars represent light or dark, respectively. (B) Representative wheel-running activity actograms of control (Co) and SCN*^Per1^*^nKo^ (top two rows), and in LHb*^Per1^*^nKo^ mice (bottom two rows). A 15-minute light pulse (LP, yellow asterisks) was applied at circadian time (CT) 10 (first column), CT14 (middle column), and CT22 (third column). The blue lines indicate onset of wheel-running activity before the LP and the red lines onset of activity after the LP. (C) Quantification of the distances between the blue and red lines in A representing the amount of phase shift in minutes after the LP at CT10 (left), CT14 (middle), and CT22 (right). SCN*^Per1^*^nKo^ significantly increases phase delays (yellow column, middle panel) and decreases phase advances (yellow column, right panel) compared to controls and LHb*^Per1^*^nKo^. Data are represented as mean ± SEM, n = 10-12, one-way ANOVA with Tukey’s multiple comparison test, *p < 0.05, **p < 0.01, ***p < 0.001, ****p < 0.0001.

Together, these results demonstrate that *Per1* expression in the SCN is required for normal light-induced phase advances and influences the magnitude of phase delays, whereas *Per1* in LHb neurons does not contribute to circadian phase shifting.

### Light-induced improvements in despair-like behavior require *Per1* in LHb neurons but not in SCN neurons

To determine how light affects despair-like behavior in mice lacking *Per1* in SCN or LHb neurons, we performed the tail suspension test (TST) at ZT6 on the third day after animals received a light pulse (LP) or no LP at ZT22. After roughly one week of recovery period, brains were collected at ZT12 to verify viral injection sites and confirm *Per1* deletion in the SCN or LHb (Fig 3A).

**Fig 3.**
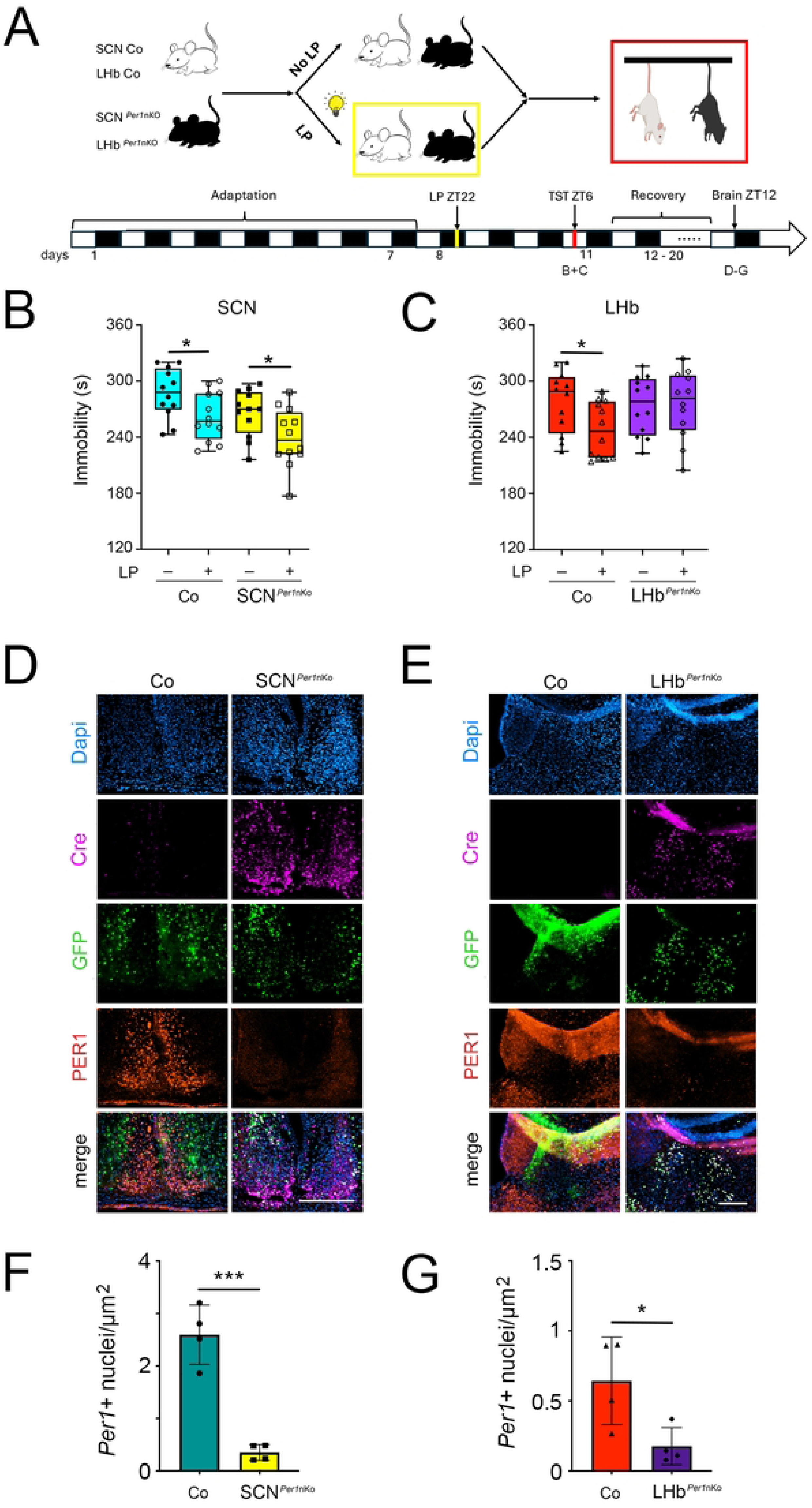
*Per1* deletion in neurons of the LHb but not the SCN abolishes the beneficial effects of light on despair-like behavior. (A) Mice were subjected to the tail-suspension test (TST) and their time of immobility was measured as a proxy for despair-like behavior. The schematic for the experimental protocol is shown. (B) A 30 min LP at ZT22 reduced immobility in the TST of SCN control (Co) and SCN*^Per1^*^nKo^ mice. Data are represented as mean ± SEM, n = 12, two-way ANOVA with Sidak’s multiple comparison test, *p < 0.05. (C) A 30 min. LP reduced immobility in the TST of LHb control (Co) but not LHb*^Per1^*^nKo^ mice. Data are represented as mean ± SEM, n = 12, two-way ANOVA with Sidak’s multiple comparison test, *p < 0.05. (D) Immunohistochemistry of SCN tissue (Dapi staining, blue) showing the expression of Cre-recombinase (pink), GFP (green), and PER1 (red) of mice receiving the control (Co) or *Per1*nKo construct. In SCN*^Per1^*^nKO^ animals Cre is present and PER1 absent in contrast to control animals. Scale bar: 200 µm. (E) Immunohistochemistry of LHb tissue (Dapi staining, blue) showing the expression of Cre-recombinase (pink), GFP (green), and PER1 (red) of mice receiving the control (Co) or *Per1*nKo construct. In LHb*^Per1^*^nKO^ animals Cre is present and PER1 absent in contrast to control animals. Scale bar: 200 µm. (F) Quantification of number of *Per1* positive nuclei per μm^2^ in the SCN of control and SCN*^Per1^*^nKO^ animals. Data are represented as mean ± SEM, n = 4, unpaired t-test, ***p < 0.001. F-test indicates similar variance (p > 0.05). (G) Quantification of number of *Per1* positive nuclei per μm^2^ in the LHb of control and SCN*^Per1^*^nKO^ animals. Data are represented as mean ± SEM, n = 4, unpaired t-test, *p < 0.05. F-test indicates similar variance (p > 0.05).

In SCN control (Co) mice (Fig. 3B, green bars) as well as in SCN*^Per1^*^nKo^ mice (Fig 3B, yellow bars), an LP at ZT22 significantly reduced immobility in the TST, consistent with previous findings in the forced swim test (FST) [16]. Loss of *Per1* in the SCN did not affect this light-induced reduction in immobility (Fig 3B, yellow bars). However, SCN*^Per1^*^nKo^ mice displayed significantly lower baseline immobility even without an LP (Fig 3B), suggesting that *Per1* in the SCN modulates the baseline behavioral state but is not required for the behavioral response to light.

In LHb control mice an LP at ZT22 also significantly reduced immobility (Fig 3C, red bars). In contrast, LHb*^Per1^*^nKo^ animals showed immobility levels comparable to untreated controls, regardless of light exposure (Fig 3C, purple bars). This is consistent with our previous findings showing that *Per1* in the LHb is required for light-mediated reduction in immobility in the FST [16]. Thus, *Per1* in LHb neurons is essential for the beneficial behavioral effects of light, whereas *Per1* in SCN neurons slightly reduces baseline immobility.

To confirm accurate viral targeting, we performed immunohistochemistry on brains from the behavioural experiments. Cell nuclei are shown in blue (Dapi staining). GFP staining (green) verified viral presence in the SCN (Fig 3D) or LHb (Fig 3E). We then assessed expression of Cre-recombinase (pink), GFP (green), and PER1 (red) (Fig 3E and 3F). Control animals showed GFP and PER1 expression but no Cre, confirming successful control virus delivery without *Per1* deletion (Fig 3D and 3E, left panels). In contrast, Cre delivery to the SCN (Fig 3D) or LHb (Fig 3E) eliminated PER1 expression in the targeted region as validated by quantification in the SCN (Fig 3F) and LHb (Fig 3G). These results confirm successful viral targeting and *Per1* deletion, supporting the conclusion that the observed behavioural phenotypes arise from region-specific loss of PER1.

### Distinct light-mediated gene expression in the SCN and LHb

To determine whether the SCN- and LHb-specific behavioral effects of light reflect distinct transcriptional responses, we performed quantitative real-time PCR analysis on SCN and LHb tissue from *Per1^fl/fl^* control, SCN*^Per1nKo^* and LHb*^Per1nKo^* mice receiving a light pulse at ZT22.

In *Per1^fl/fl^* control animals, *Per1* expression increased after the LP in the SCN (Fig 4A, top left and middle panels, blue bars), consistent with previous reports [7, 23]. Because *Per1* contains two promoters with alternative first exons (*Per1_p1* and *Per1_p2*; [24]), we examined promoter-specific induction. *Per1_p1* was induced by light in both the SCN and LHb of *Per1^fl/fl^* mice (Fig 4A, left panel, blue bars) and in all other genotypes except those lacking *Per1* in the respective region (Fig 4A, left panel, yellow and purple bar). In contrast, *Per1_p2* was induced by light only in the SCN and only in genotypes with intact *Per1* expression (Fig 4A, middle panel, top, yellow bar). No *Per1_p2* induction was observed in the LHb for any genotype (Fig 4A, middle panel, bottom). Thus, *Per1_p2* activation is SCN-specific. As previously reported [7, 16, 23], *Per2* expression was not significantly affected by light at ZT22 in either region across all genotypes.

**Fig 4.**
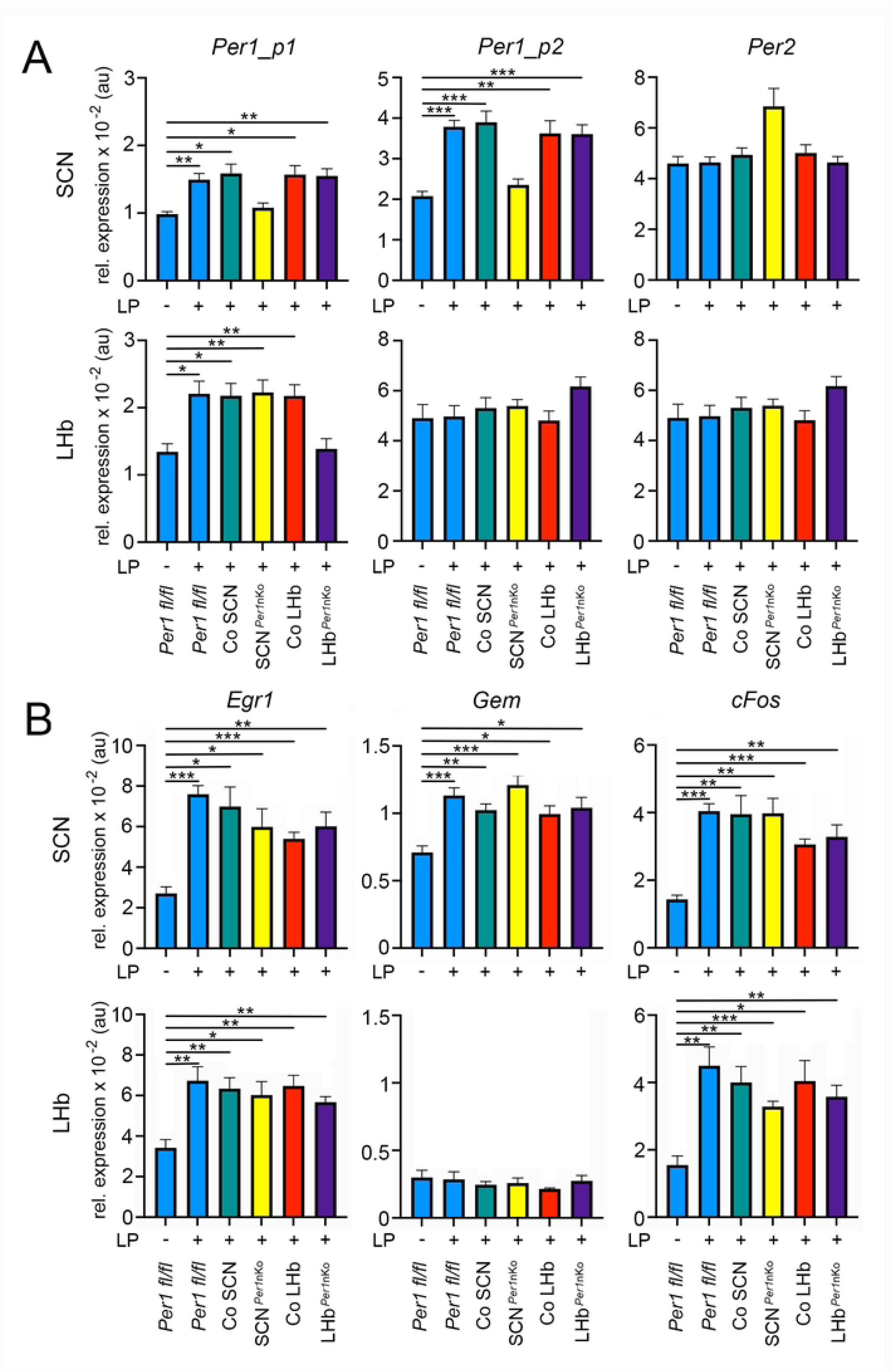
Gene expression in the SCN and LHb of mice after a light pulse at ZT22 compared to control animals. (A) *Per1* and *Per2* gene expression in SCN (top row) and LHb (bottom row). *Per1* gene expression was measured from promoter 1 for exon 1A (*Per1_p1*) and promoter 2 for exon 1B (*Per1_p2*) of the *Per1* gene. *Per1* was light inducible from both promoters in the SCN but only from promoter 1 in the LHb. *Per2* was not significantly induced by light at ZT22. Data are presented as mean ± SEM, n = 8, Brown-Forsythe and Welch ANOVA test with Dunnett’s T3 multiple comparisons test, * p < 0.05, ** p < 0.01, *** p < 0.001. (B) *Egr1, Gem,* and *cFos* gene expression in SCN (top row) and LHb (bottom row). *Egr1* and *cFos* were both light inducible in both the SCN and LHb, whereas *Gem* was only inducible in the SCN and not in the LHb. Data are presented as mean ± SEM, n = 8, Brown-Forsythe and Welch ANOVA test with Dunnett’s T3 multiple comparisons test, * p < 0.05, ** p < 0.01, *** p < 0.001.

We next examined additional light-inducible genes, including immediate early genes and other clock genes [25–28]. The immediate early genes *Egr1* and *cFos*, were robustly induced by light in both SCN and LHb tissue of all genotypes, including *Per1* knockouts (Fig 4B, left and right panels). This indicates that their induction is *Per1*-independent and not region-specific. In contrast, *Gem,* a regulator of plasma membrane signaling [29], was induced by light exclusively in the SCN across all genotypes (Fig 4B, middle panels), demonstrating region-specific induction that does not require *Per1*. The clock genes *Dec1* and *Bmal1* were not light-inducible at ZT22 (S1A Fig), consistent with previous reports [30, 31]. Interestingly, however, we observed a reduction of *Bmal1* expression in animals lacking *Per1*. The exclusive expression of *Irs4* in the SCN and *Gpr151* in the LHb showed that the tissues used for qPCR analysis were correctly harvested (S1B Fig). The expression of *eGfp* showed correct injection of viral particles into the SCN and LHb (S1B Fig).

Together, these findings show that immediate early genes are induced by light at ZT22 independently of *Per1* and independently of brain region. In contrast, *Per1* itself shows promoter-specific and region-specific induction: promoter 1 is light-responsive in both SCN and LHb, whereas promoter 2 is light-responsive only in the SCN. These results support the hypothesis that the SCN and LHb interpret light signals through distinct *Per1*-dependent pathways that differentially regulate circadian and mood-related behaviors.

## Discussion

The present study shows that light regulates circadian phase and despair-like behavior through two anatomically and distinct *Per1*-dependent mechanisms. By selectively deleting *Per1* in either the SCN or the LHb, we provide evidence that the circadian and affective effects of light can be dissociated at the expression levels of transcripts emerging from either of the two promoters. This finding challenges the long-standing assumption that light’s antidepressant-like effects are secondary to its circadian phase-shifting properties and instead supports a model in which mood regulation and circadian entrainment represent parallel outputs of photic signaling (Fig 5).

**Fig 5.**
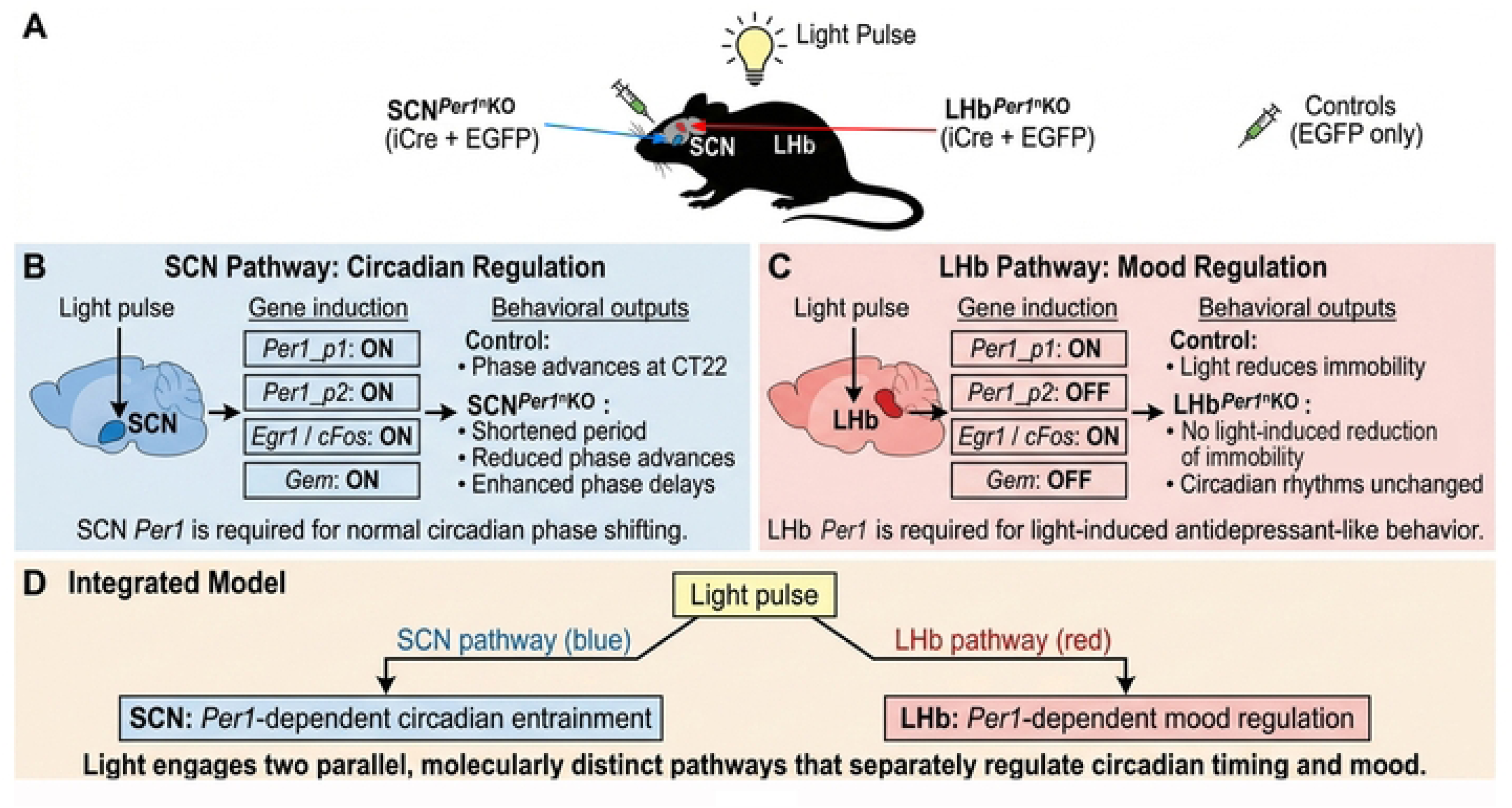
Distinct *Per1*-dependent light-responsive pathways in SCN and LHb. (A) Targeted *Per1* deletion in SCN or LHb was combined with a light pulse at ZT22. (B) In the SCN, light activates *Per1_p1/p2*, *Egr1/cFos*, and *Gem*, enabling normal circadian phase shifting; SCN*^Per1^*^nKO^ mice show impaired light-induced entrainment. (C) In the LHb, light induces *Per1_p1* and *Egr1/cFos* but not *Per1_p2* or *Gem*; LHb*^Per1^*^nKO^ mice lose light-induced antidepressant-like behavior while maintaining normal circadian rhythms. (D) Model: light engages parallel, molecularly distinct *Per1*-dependent pathways that separately regulate circadian timing (SCN) and mood (LHb). (Figure made with Copilot and Gemini AI-agents).

Consistent with classical phase response curves [11], light at CT22 induced robust phase advances in control mice (Fig 2). However, these advances were markedly reduced in SCN*^Per1^*^nKO^ animals, confirming that *Per1* expression in the SCN neurons is essential for normal photic entrainment [22, 32]. This aligns with earlier work showing that *Per1* induction is a key early event in SCN phase resetting [7, 8, 33]. The larger phase delays observed at CT14 in SCN*^Per1^*^nKO^ mice further suggest that *Per1* contributes also to shaping the magnitude of delays, possibly by modulating the balance between *Per1*-driven and *Per2*-driven resetting pathways [9, 10, 32, 34]. In contrast, LHb*^Per1nKo^* mice displayed normal phase shifts at all circadian times, demonstrating that *Per1* in the LHb is not required for circadian entrainment (Fig 2). This is consistent with the LHb’s limited role in circadian pacemaking and supports the idea that LHb photic responses serve non-circadian functions [13, 35].

A central finding of this study is that the antidespair-like effect of light at ZT22 is abolished in LHb*^Per1nKo^* mice but preserved in SCN*^Per1^*^nKO^ animals (Fig 3B and 3C). This indicates that *Per1* in LHb neurons is necessary for the behavioral response to light in the TST, whereas SCN *Per1* is dispensable. The LHb is a key regulator of aversive learning [36], negative reward prediction [37, 38] and despair-like behavior, and its hyperactivity is associated with depressive-like phenotypes [39, 40]. Inhibition of the LHb via activation of the ventral lateral geniculate nucleus and intergeniculate leaflet (vLGN/IGL) projecting neurons reversed the depressive-like phenotypes of mice exposed to a battery of aversive stimuli [13]. Similar behavioural observations have been made in response to aversive treatment, following administration of bright light therapy [41]. Thus, light-induced *Per1* expression may therefore modulate LHb excitability or downstream monoaminergic circuits, producing the observed reduction in immobility in the TST. Interestingly, SCN*^Per1^*^nKO^ mice exhibited reduced baseline immobility even without light exposure (Fig 3B). This suggests that SCN *Per1* contributes to the regulation of baseline behavioral state – potentially through altered circadian parameters (Fig 1C), sleep architecture, or hormonal rhythms – but does not mediate the acute behavioral effects of light in the TST.

qPCR analyses revealed that the SCN and LHb interpret light through different promoter-specific and gene-specific transcriptional programs. *Per1* promoter 1 was light-responsive in both brain regions, whereas promoter 2 was activated only in the SCN (Fig 4A), suggesting that SCN-specific transcription factors or chromatin states enable promoter 2 induction. Immediate early genes, such as *Egr1* and *cFos*, were induced in both regions independently of *Per1* (Fig 4B, left and right panels), indicating that early photic signalling is intact in knock-out animals and that *Per1* acts downstream of initial light detection. The SCN-specific induction of Gem further highlights divergent transcriptional repertoires (Fig 4B, middle panels). Together, these data support a model in which the SCN and LHb receive similar photic input but decode it through distinct molecular pathways, enabling light to simultaneously influence circadian timing and mood via separate mechanisms.

These findings have direct translational relevance. Light therapy is widely used to treat seasonal affective disorder (SAD) and is increasingly explored for major depressive disorder (MDD) [18, 42, 43]. The prevailing assumption has been that mood improvements arise from correcting circadian misalignment. Our results challenge this view by showing that light’s antidepressant-like effects do not require circadian phase shifting and instead depend on *Per1* signaling in the LHb. This dissociation suggests that therapeutic strategies could be optimized by targeting LHb-specific photic pathways, potentially through wavelength-specific stimulation, targeted neuromodulation (e.g. deep brain stimulation) or pharmacological enhancement of *Per1*-related signaling, without necessarily altering circadian rhythms. Such approaches may benefit patients who are sensitive to circadian disruption or who require rapid mood improvement.

Several questions remain open. First, the downstream targets of *Per1* in LHb neurons remain unknown. Transcriptomic or single-cell approaches could identify candidate genes mediating the behavioral response. Second, the circuit mechanisms linking LHb *Per1* induction to reduced despair-like behavior warrant investigation, particularly interactions with the ventral tegmental area, dorsal raphe, and rostromedial tegmental nucleus. Third, it will be important to determine whether the effects of light through *Per1* in the LHb involve a functional clock or whether this signaling pathway is clock independent. Finally, whether similar dissociations exist in humans remains an open question. Functional imaging studies combined with genetic or chronotype-based stratification may help determine whether LHb-specific photic responses contribute to the therapeutic effects of light in mood disorders. Such a dissociation likely exists in humans, as deep brain stimulation of the lateral habenula has been associated with remission of major depression [44, 45] and light treatment could be viewed as a form of deep brain stimulation.

## Methods

### Mouse strains used and their housing conditions

Male and female (50:50) *Per1 fl/fl* mice (described in [16]; EMMA: 14846) aged 2 months were placed in isolated light cabinets in type II L cages. Entrainment was done in a constant 12:12 h light /dark cycle. Temperature (22 ± 2 °C monitored by temperature sensor, Technoline WS-9410, Berlin, Germany), humidity (40–50% monitored by humidity sensor, Technoline WS-9410, Berlin, Germany), and illumination (1000 Lux monitored by Luxmeter, Testo, GmbH & Co, Titisee-Neustadt, Germany) were kept constant in all cabinets. For the wheel-running experiments mice were housed individually in cages (L: 280 mm × W: 105 mm × H: 125 mm) containing a running wheel made of steel (115 mm in diameter, Trixie GmbH, Tarp, Germany). All mice were provided with sufficient woodchip bedding as to not block the running wheel, and enrichement materials such as: red-square house, piece of carton, neslet (5x5 cm), and an open-sided tube. Food and water were provided *ad libitum.* All experiments and procedures were performed according to the Schweizer Tierschutzgesetz guidelines and approved by the Canton of Fribourg and the cantonal commission for animal experiments (2023-08-FR, 35721).

### Stereotaxic surgeries

The adeno-associated viruses (AAV) were provided by the Viral Vector Facility (VVF) of the Neuroscience Center Zürich (ZNZ). For injection into the SCN of *Per1* fl/fl, the following virus was used as a control: ssAAV-9/2-hSyn1-EGFP-WPRE-hGHp(A) with a titer of 2.9 x 10^13^ viral genome equivalent/ml, and for Cre delivery the ssAAV-9/2-hSyn1-EGFP_iCre-WPRE-hGHp(A) with a titer of 8.5 x 10^12^ viral genome equivalent/ml. For injections into the LHb, the control virus was ssAAV-6/2-hSyn1-EGFP-WPRE-hGHp(A) with a titer of 9.3 x 10^12^ viral genome equivalent/ml, while for the Cre delivery we used ssAAV-6/2-hSyn1-EGFP_iCre-WPRE-hGHp(A) with a titer of 6.8 x 10^12^ viral genome equivalent/ml. AAV vectors and plasmids required for their synthesis are available on the VVF website (https://www.vvf.uzh.ch/en.html).

2-months old mice were selected for stereotaxic injections. Mice were placed under anaesthesia using isoflurane/oxygen (2% isolflurane v/v, 1 l O_2_/min, induction: 5% v/v, 1 l O_2_/min). For analgesia, 5 mg/kg carprofen and 0.15 mg/kg buprenoprhrine were administered 30 minutes before the procedure and 2% lidocain was applied locally during the surgery. Eyes were protected using a Vitamin A retinoli palmitas (Bausch & Lomb Swiss AG, Zug, Switzerland). The head of the mouse was fixed in the stereotaxic instrument (Neurostar, Tubingen, Germany, Serial Number: SD877), and fitted with the anaesthesia inhalation mask of the stereotaxic instrument. The bregma and lambda were used as refference points to corrrect for the tilt and scalling of each individual’s brain. Following these corrections, a pulled glass pipette (Wiretrol II, Drumont Scientific, Bromall, PA, USA) was used to inject 100 nl (50nl into each hemisphere) of the AAV viruses into the SCN, or 200 nl (100 nl into each hemisphere) into the LHb. The specific coordinates for the SCN were AP: -0.46 mm, ML: +/- 0.16 mm, DV: 5.62 mm, angle 0^0^, and for the LHb: AP: -1.82 mm, ML: +/- 0.49 mm, DV: 2.58 mm, angle 0^0^. Following the injection of the AAV virus, the glass pippette was held into place for 5 mins, after which it was raised by 0.1mm and held into place for another 3 mins to prevent the upward spread of the virus. At the end of the procedure, the skin of the mouse above the skull was sealed using GLUture (Zoetis, Kalamazoo, MI, USA). Mice were allowed to recover for 3 weeks before being subjected to any behavioral tests.

### Light Pulse for Wheel Running Experiments (Aschoff Type I Protocol)

The experimental set-up was designed and performed according to the Aschoff Type I protocol [46, 47]. Mice were entrained to a 12:12 light:dark (LD) cycle for 2 weeks, then released into constant darkness (DD) for 2 weeks using an automatic digital timer. To calculate the LP, the final 10 days of wheel-running activity in DD were analyzed in ClockLab3 (Acquisition v3.208; Analysis v6.0.36, Lafayette, IN, USA) software, to determine each mouse’s individual period length and predict activity onset at circadian time (CT) 12 the following day. A 15-minute LP (1000 LUX) was then administered at CT10 (or CT14, CT22 for subsequent pulses) in a separate cabinet, after which mice were returned to their original cabinet for another 2 weeks of free-running recording. After the final DD phase, mice were re-entrained to 12:12 LD for 2 weeks.

### Monitoring of Circadian Locomotor Activity Rhythm

Circadian locomotor activity was quantified by monitoring running-wheel revolutions with a vertically mounted magnetic sensor attached to the wheel axis outside the cage. Wheel turns were recorded in 1-min bins using the ClockLab 3 data acquisition and analysis software, as detailed previously [48, 49].

### Analysis of Circadian Locomotor Activity Rhythm Parameters

Phase shifts (i.e., phase resetting) in constant darkness (DD) following each light pulse (LP), along with other circadian locomotor activity rhythm parameters (e.g., actograms, period length, amplitude, total locomotor activity counts, and relative rhythm power via FFT), were analyzed using ClockLab software. Phase shift magnitude for each LP was determined by drawing a regression line through 10 consecutive activity onsets prior to the pulse, and a second line through 10 activity onsets following the LP (excluding the first two transitional days). The difference between these lines on the day of the LP onset represents the phase shift value.

### Light Pulse for the Behavioral Test of Tail Suspension

Mice were exposed to polychromatic ligth at 1000 lux in their home cages for 30 minutes at ZT22 (i. e., until ZT22.5). The control group of mice was handled and treated in the same way (cage movement, presence of the experimenter) in the dark, receiving no light pulse. *Tail Suspension Test (TST)* was performed at ZT6. Up to 4 mice were tested at the same time in suspension boxes (55cm high, 15 cm in diameter), designed in our department. Each compartment ensured there was sufficient space for the mouse to undergo the test, without touching the walls. Their tails were suspended on an aluminium bar sitting on top of the compartment by a 17cm tape (allowing 2 extra cm to wrap around the tails). Small tubes were placed at the start of their tails as to stop the mice from climbing onto them. All mice performed a 6-minute session, while the experimenter left the room to minimise disturbance. The sessions were recorded with a video camera (Sony HDR-CX450), and the videos were scored manually. The output of the test was expressed as immobility time recorded during the entire duration of the 6 minute session. Frontal paw movements, and swinging of the mice from previous movement bouts were not counted as mobility. Scoring was performed blinded with the assessor not knowing the genotype or treatment of the mice.

### Tissue collections and Immunohistochemistry

Mice were perfused at ZT12 with 4% paraformaldehyde (PFA). Brains were carefully isolated to preserve the optic nerve region and hence the suprachiasmatic nucleus (SCN), then post-fixed in 4% PFA for 12–15 hours before cryoprotection in 30% sucrose. SCN and LHb cryosections (40 μm) from each brain were collected into 24-well plates.

For surgery target validation, sections were counterstained with DAPI (Roche; 1:5000 in 1× PBS/0.2% Triton X-100) for 20 minutes, followed by 2 washes in 1× PBS/0.2% Triton X-100 before mounting on glass slides. Sections were imaged using the Hamamatsu NanoZoomer S60 (Solothurn, Switzerland).

For antibody immunohistrochemistry stainings, sections underwent 20 minutes of washing in 1× PBS, followed by three 10-minute washes in 1× PBS with 0.2% Triton X-100. The sections were then treated for 15 minutes in 1× PBS with 1% Triton X-100, followed by three additional washes in 1× PBS/0.2% Triton X-100. Sections were then blocked for 3 hours at room temperature in 1× PBS/0.2% Triton X-100 containing 10% donkey serum (Sigma, Catalog #D9663).

Primary antibodies targeting PER1, and Cre (Table 1), diluted in blocking solution, were applied overnight for 2 days at 4°C. Sections were washed five times (10 minutes each) in 1× PBS/0.2% Triton X-100, then incubated for 2 hours at room temperature with secondary antibodies (Table 1) diluted in blocking solution. This was followed by five 10-minute washes in 1× PBS/0.2% Triton X-100 and an overnight wash at 4°C. Lastly sections were counterstained with DAPI (Roche; 1:5000 in 1× PBS/0.2% Triton X-100) for 20 minutes.

**Table 1.** Antibodies.

| **Antibody** | **Species** | **Company** | **Catalog number** | **Lot Number** | **Dilution** |
| --- | --- | --- | --- | --- | --- |
| Anti-mPER1<br>(residues 6-21) | Rabbit | Merck Millipore | AB2201 | 3480987 | 1:8000 |
| Anti-Cre<br>Recombinase,<br>clone 2D8 | Mouse | Merck Millipore | MAB3120 | 4206901 | 1:1000 |
| DyLight™ 405-AffiniPure<br>Donkey Anti-Mouse IgG<br>(H+L) | Donkey | Jackson<br>Immunoresearch | 715-475-150 | 168170 | 1:1000 |
| Donkey Anti-Rabbit IgG H&L<br>(Alexa Fluor<br>568) | Donkey | Abcam | Ab175693 | GR3264869-3 | 1:1000 |

Fluorescent Z-stack images were acquired using the Leica DM6B microscope (Leica Microsystems, Heerbrugg, Switzerland). Images were processed in Imaris software (version 11.0.1, Oxford Instruments, Buckinghamshire, UK), where deconvolution and background substraction were performed. Nuclei quantification was performed using the spots function of the software.

### Quantitative polymerase chain reaction (qPCR) analysis

RNA from SCN or LHb tissue was isolated with Nucleo Spin RNA XS (Macherey-Nagel, Oensingen, Switzerland) according to manufacturer’s instructions. Reverse transcription starting from 0.5 mg of totalRNA was performed using the SuperScript IV VILO master mix (Thermo Fisher Scientific, Vilnius, Lithuania). After reverse transcription, samples were diluted 40 times. 5μl of the dilued sample was added to 7.5 μl of KAPA probe fast universal master mix (Merck, Darmstadt, Germany) and 2.5 μl of primers/probe mix (listed in Table 2) (150 nM of primers, 33.3 nM TaqMan probe). Samples were run on a Rotor-Gene RG-6000 qPCR machine equipped with a Rotor-disc 100 rotor (Qiagen, Hilden, Germany). All targets of interest were normalised against the four housekeeeping genes *Nono*, *Gapdh*.

**Table 2.**
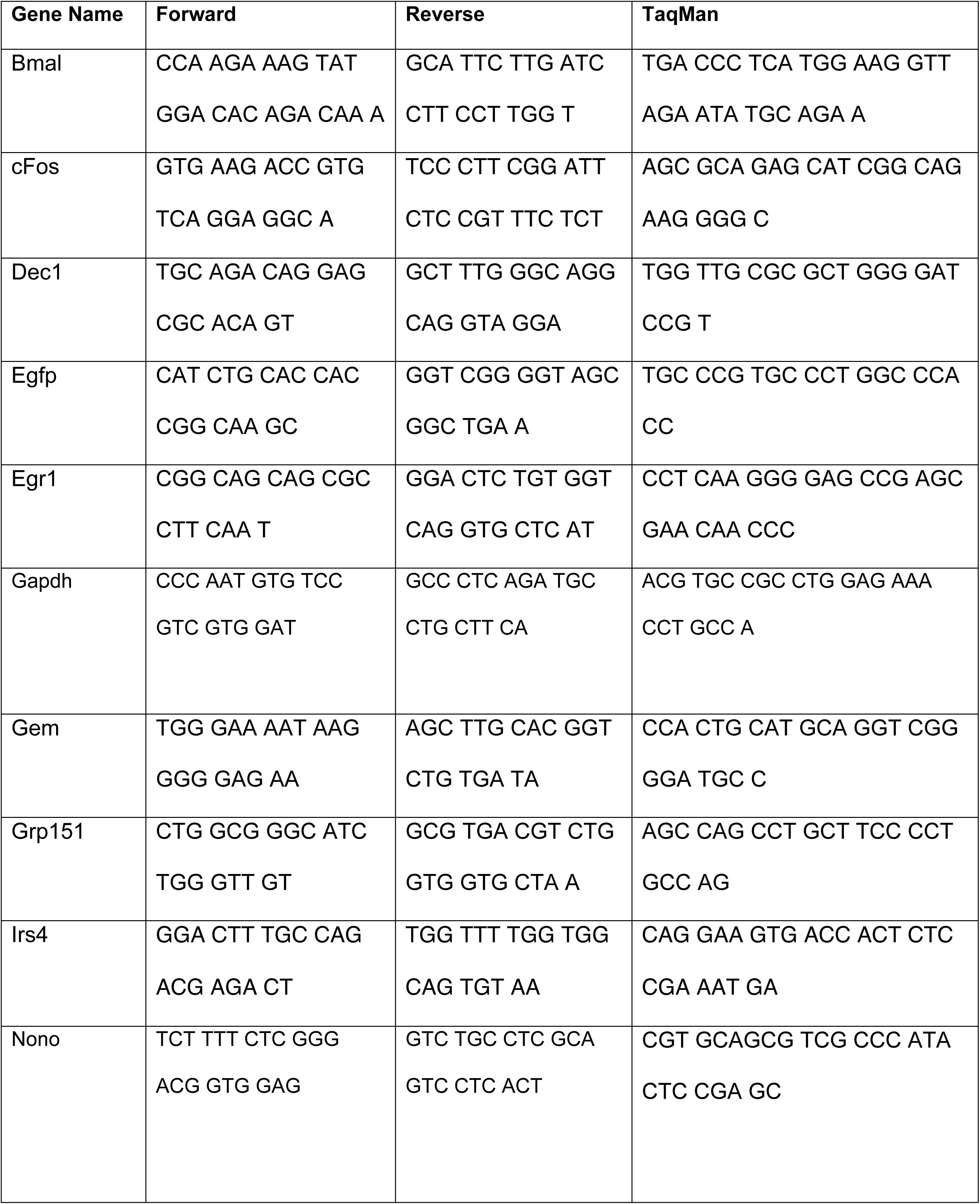
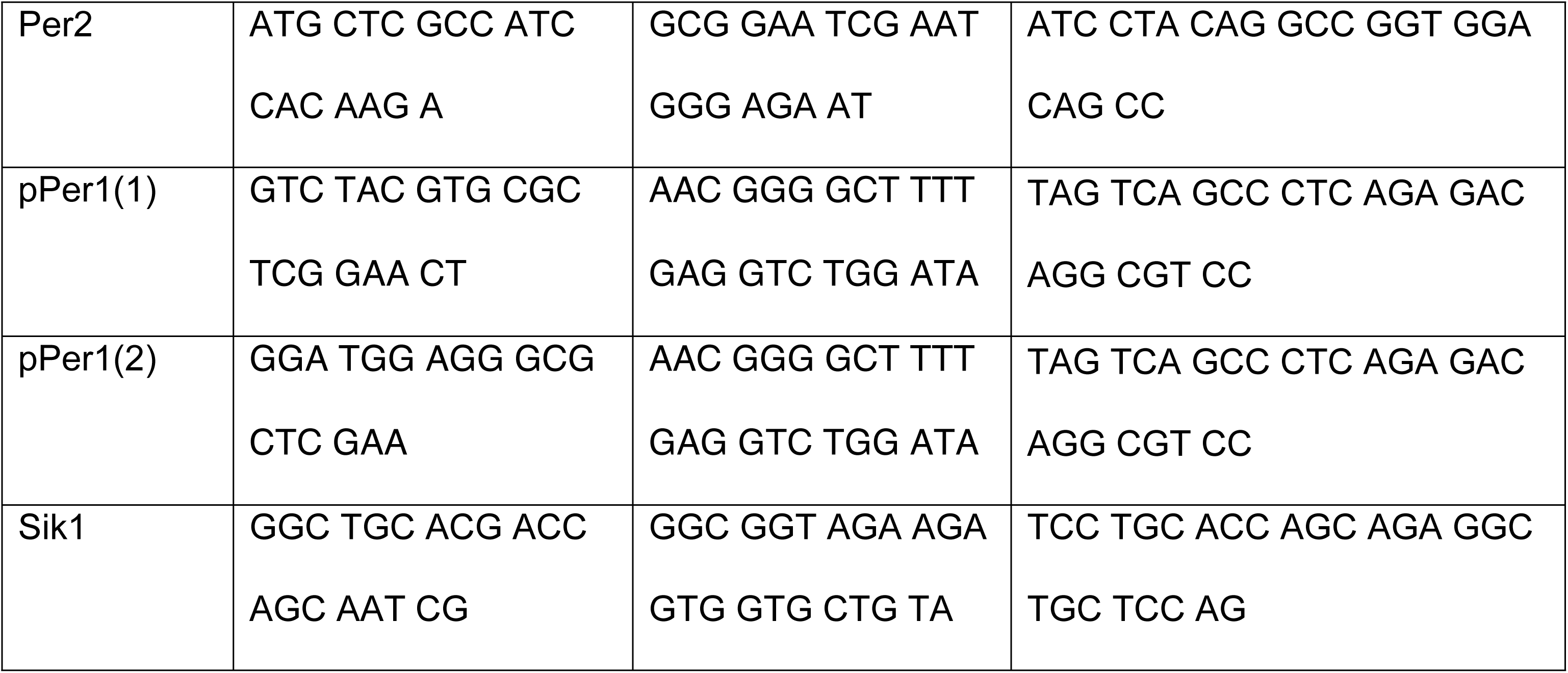
PCR-primers.

### Statistical Analyses

According to the experimental designed, the statistical tests of two-way ANOVA, followed Sidak’s multiple comparison test, or one-way ANOVA followed by Tukey’s or Dunnett’s multiple comparison tests were used. The statistical tests were performed using GraphPad Prism (version 10.4.1). Data are considered significant when the p value < 0.05.

## Supporting Information

**S1 Fig. Gene expression in the SCN and LHb of mice after a light pulse at ZT22 compared to control animals.** (A) *Dec1*and *Bmal1* gene expression is shown in the SCN (top row) and LHb (bottom row). *Dec1* and *Bmal1* are not induced by light. *Bmal1* expression was reduced in the tissues lacking *Per1*. (B) Expression of control genes are shown. The SCN specific *Irs4* gene is only expressed in the SCN but not the LHb (left panels), whereas the LHb specific gene *Gpr151* is only expressed in the LHb but not the SCN (middle panels), indicating correct harvesting of tissues used for qPCR analysis. The *Gfp* expression (right panels) is only present in the tissue transfected with the viral constructs. Data are presented as mean ± SEM, n = 8, Brown-Forsythe and Welch ANOVA test with Dunnett’s T3 multiple comparisons test, * p < 0.05, ** p < 0.01, *** p < 0.001.

## Acknowledgments

We thank Antoinette Hayoz, Maude Marmy, Ilona Bouhier and the Bioimage platform (University of Fribourg) for technical support.

